# Prevalence of HIV-1 infection among foreign applicant to residency in Shanghai, China, 2005-2016

**DOI:** 10.1101/560854

**Authors:** Jia Qin, Ru Zhou, Jing Xia, Weixin Wang, Jun Pan, Jiahong Pan, Xuan Zhou, Qi Zhang

## Abstract

**Background:** Shanghai is one of the biggest cities which have the highest number of entry travelers from all over the world. The HIV(human immunodeficiency virus) infection status of this population can reflect global trends of HIV prevalence to a certain extent.

**Methods:** A retrospective cohort study was conducted to reveal the prevalence and characteristics of HIV-1 infection among entry travelers who applied to residency in Shanghai. The HIV-1 infection rate was estimated based on the detection of HIV-1 antibody.

**Results:** Among 50830 entry travelers who applied to residency in Shanghai(2005-2016), 245 were determined HIV-1 positive with an infection rate of 0.48%. The detection rate of HIV was significantly higher in male (*P*<0.0001). Those aged 18-30 years, 31-40 years and >40years accounted for 34.3%,39.6% and 26.1% respectively of the infected population. Although there was no trend of increase in HIV-1 prevalence rates (Cochran-Armitage Z =2.543, P =0.111),proportions of individuals infected through homosexual transmission increased over the study period (Cochran-Armitage Z =5.41, P<0.001), while the proportions infected through heterosexual(Cochran-Armitage Z=3.38, P=0.001).

**Conclusion:** The rate and characteristics of HIV-1 infection among foreign applicant to residency in Shanghai were revealed in the study. The results could provide the necessary epidemiological data for monitoring the HIV-1 epidemic among entry international travelers and to further contribute to the establishment of relevant policies and regulations for HIV control and prevention.

## Background

Since the AIDS epidemic peaked in 1993, it has become the leading cause of death among 25 to44-year-old individuals, and the eighth most common cause of death worldwide ^[1]^. Although human immunodeficiency virus (HIV)-1 infection and mortality rates have been declining because of the highly active antiretroviral therapy and effective prevention measures globally, an estimated 35.3 million people are living with HIV worldwide with approximately 2.3 million new infections per year. HIV infections are considered to be the global threat of present era ^[2,3]^.There is still urgency in expanding HIV testing and treatment across the world ^[4]^.The international health regulations (2005) entered into force in June 15th, 2007. The new regulations fully respect the sovereignty of the State Party and the human rights of travelers, emphasizing the involuntary physical examination of travelers, and the absence of a certain physical examination as a condition for allowing travelers to enter. China’s State Council promulgated and implemented the “AIDS prevention and control regulations” in March 1st, 2006, to clearify the work requirements and tasks of inspection and quarantine system. In May 30th, 2007, AQSIQ examined and approved the “port AIDS prevention and control measures”, the port entry&exit personnel AIDS quarantine, monitoring, prevention and control and financial security has specific provisions and requirements. The implementation of these policies provides an opportunity for the inspection and quarantine system to strengthen the intervention of AIDS, and a strong support for the development of AIDS prevention and control work for Entry-Exit personnel. On April 24 of 2010, the executive meetings of the State Council adopted *“*the Decision of the State Council on Amending *‘*Detailed Rules for the Implementation of the Law of the People’s Republic of China on Frontier Health Quarantine Inspection’”, which clearly stipulated that the entry of foreign AIDS patients was no longer restricted. This regulation is consistent with most developed countries. Shanghai is a major destination for international business and trade and thus has a large population of cross-border travelers. Entry travelers and their activities often lead to the spread of infectious diseases in China and even on a global scale, and therefore these people are often considered a bridge population for infectious disease transmission ^[5]^. Furthermore, such a large bridge population could promote HIV cross-border transmission, leading to a higher HIV prevalence in the world. Strengthening surveillance and attention to this group is crucial to the prevention of HIV/AIDS, particularly in the border areas which are filled with business people and tourists. In addition, the investigation of changes in HIV-1 infection and the associated risk factors over time could be important for the prevention of HIV-1 in this risk group. Although HIV infection among cross-border travelers has been reported in other major cities ^[2,6]^, it is still unclear in Shanghai. Therefore, to investigate the HIV epidemic characteristics among entry travelers, a large-scale retrospective study of HIV-1 prevalence was conducted on cross-border travelers who entered China through Shanghai ports from 2005-2016. The objective of this research was to reveal the epidemiological characteristics of HIV-1 infection among international entry travelers in Shanghai to help policy makers identify and address the specific needs for HIV prevention, control and treatment.

## Methods

### Study population and data collection

From January 2005 to December 2016, there were approximately 5000 entry travelers on average in Shanghai annually. Those foreign applicants who apply for staying in China for one or more years need to undergo physical examinations in Shanghai International Travel Healthcare Center, which is affiliated to Shanghai Entry-Exit Inspection and Quarantine Bureau and is responsible for the health assessment and epidemiological surveillance of people entering and exiting Shanghai. Routine examinations include full blood count, serum biochemistry tests, and the detections of major infectious diseases. As part of the infectious disease package, it is mandatory for all applicants to undergo the HIV test, and the patients who had physical examination reports done elsewhere could be asked to repeat the test depending on date and antibody status of the last available test. Before the entry, the applicant who had voluntarily declared being HIV positive had to be re-examined to confirm the diagnosis. Initial screening was carried out with an enzyme-linked immunoassay (ELISA) or an enzyme immunoassay (EIA). The same assay was repeated for positive samples. If two of three test results were positive, the sample was subject to further FDA-approved supplemental/confirmatory assays, such as HIV-1 Western Blot (Genetic Systems HIV-1 Western Blot, Bio-Rad Laboratories, Redmond, Washington, US) and/or an immunofluorescent antibody (IFA) test (Fluorognost HIV-1 IFA, Waldheim Pharmazeutika, GmbH, Vienna, Austria). The diagnosis of HIV was based on HIV assays approved by Chinese Food and Drug Administration (FDA).Detection methods and quality control strictly followed the “National AIDS detection technical specifications” and “National AIDS detection work management measures”^[7]^.A face-to-face interview was conducted in a private room to gather information on socio-demographic characteristics for those with final diagnosis of HIV, including age, gender, nationality, travel history, occupation, education, family situation and risk behaviors of transmission. The respondents who did not complete the survey were regarded as missing data and were not included in the statistical analyses.

The nations of all HIV patients were divided into three category, namely developed, developing and underdeveloped countries, respectively. For the current 2016 fiscal year, low-income economies were defined as those with a GNI per capita, calculated using the World Bank Atlas method, of $1,045 or less in 2014; middle-income economies were those with a GNI per capita of more than $1,045 but less than $12,736; high-income economies were those with a GNI per capita of $12,736 or more; Lower-middle-income and upper-middle-income economies were separated at a GNI per capita of $4,125^[8]^.

### Data Analysis

The statistical analysis was carried out using SPSS, version 12.0 soft ware package (SPSS Inc., Chicago, IL, USA). Characteristics between the groups were compared using χ2 tests. Cochran-Mantel-Haenszel was used to compare the differences of the total population while controlling for time factors. Time trends were calculated using Cochran-Armitage trend test. The results with P values<0.05 were considered to be statistically significant.

### Ethics statement

The design of this research was reviewed by Shanghai Entry-Exit Inspection and Quarantine Bureau and approved by Shanghai Medical Ethics Committee. All personally identifiable information for the study population was stripped and the individual research identification number was assigned before the data set was sent to investigators for analyses.

## Results

### Characteristics of HIV-1 positive travelers

From 2005 to 2016, a total of 50830 travelers who entered China through Shanghai port were subject to HIV-1 infection screening test in Shanghai International Travel Healthcare Center. The annual recruited populations (range 2640-5764) were not significantly different among the sampling years. According to ELISA and WB results, 245 individuals were detected to be infected with HIV-1, resulting in an infection rate of 0.48%. Of the 245 HIV-1 infected cross-border travelers entering at Shanghai border, 215 (87.8%) were male and 30 (12.2%) were female, with a male-to-female ratio of 7.17:1. The detection rate of HIV in male was significantly higher than that in female (χ2=62.584,P<0.0001). The age range of this population was 18-69 years old (median, 34 .2 years). Those aged 18-30 years, 31-40 years and >40years accounted for 34.3%, 39.6% and 26.1% of the infected population, respectively. Most of the positive cases were employed(53.88%) and had a secondary or higher degree(81.22%). Moreover, 54.70% of the positive cases were not married and 40.82% of them were from low-income countries.

### Infection rates over the years

Infection rates over the years were listed in Table 2. Although the annual detection rates were different, the results from trend analysis showed that there was no increase in trend of HIV-1 prevalence rates among the sampling years (Cochran-Armitage Z =2.543,P =0.111) (Table 2). Trend analysis based on gender classification indicated that neither male(Z =1.944,P =0.163) nor women(Z =0.198,P =0.656)had increase trend of HIV-1 positive rates.

**Table 1.**
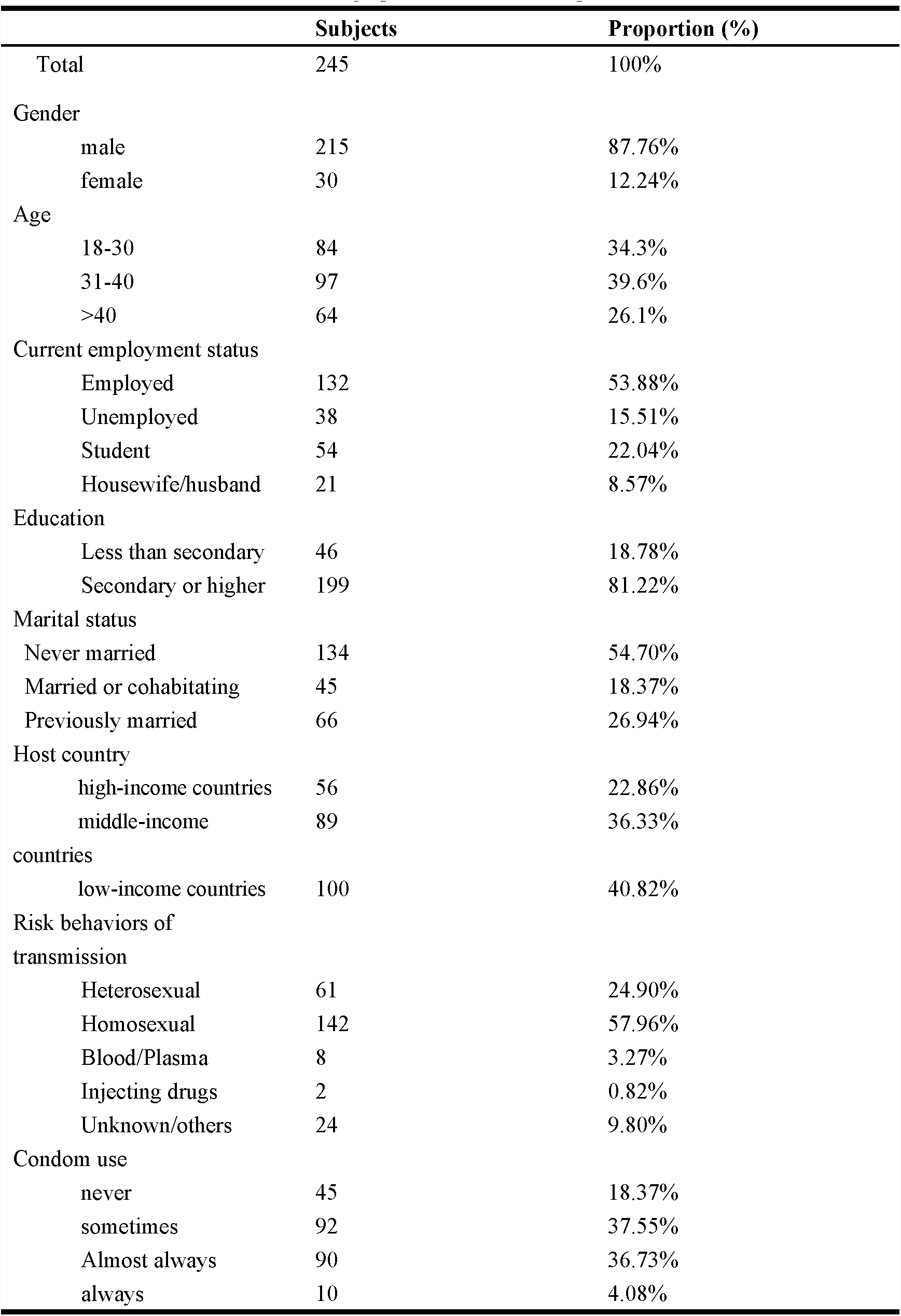
Demographic characteristic of positive cases

**Table 2.** HIV antibody positive rates (/10,000) among entry travelers 2005-2016

| Year | Male | Female | Total |
| --- | --- | --- | --- |
| 2005 | 65(10/1532) | 18(2/1108) | 45(12/2640) |
| 2006 | 54(10/1854) | 8(1/1202) | 36(11/3056) |
| 2007 | 59(13/2198) | 15(2/1374) | 15(15/3572) |
| 2008 | 58(15/2588) | 39(6/1532) | 51(21/4120) |
| 2009 | 45(12/2665) | 14(2/1445) | 34(14/4120) |
| 2010 | 80(15/1878) | 12(2/1676) | 48(17/3554) |
| 2011 | 84(19/2254) | 9(2/2136) | 48(21/4390) |
| 2012 | 85(20/2353) | 10(2/2023) | 50(22/4376) |
| 2013 | 65(17/2604) | 20(4/1969) | 46(21/4573) |
| 2014 | 99(29/2934) | 5(1/2062) | 60(30/4996) |
| 2015 | 59(27/4541) | 18(2/1128) | 51(29/5669) |
| 2016 | 77(28/3633) | 19(4/2131) | 56(32/5764) |
| Total | 69(215/31034) | 15(30/19796) | 48(245/50830) |

\* Detection rate = positive number / number of samples

### The change of transmission route

Proportions of individuals infected through homosexual transmission increased over the study period (Cochran-Armitage Z =5.41,P<0.001), while the proportion infected through heterosexual declined over time (Cochran-Armitage Z =3.38,P =0.001). The proportion infected through injecting drug also declined over time (Z=3.52,P<0.001). With the change in major HIV-1 risk factors, the heterosexual spreading of HIV-1 was gradually increasing to be the main transmission route, compared to syringe/needle sharing.

**Figure 1.**
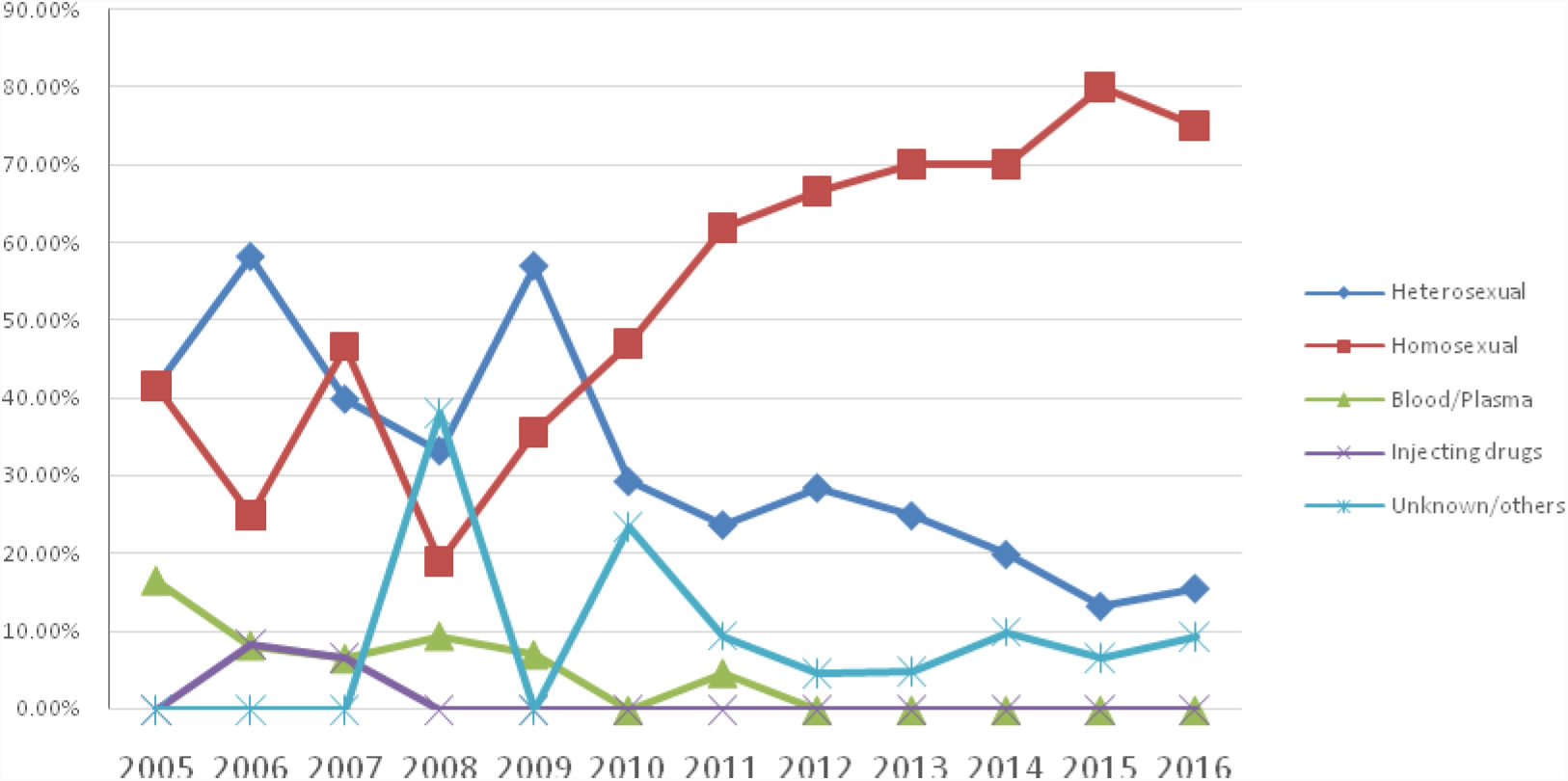
The change of transmission route over the years (Percentage of patients)

## Discussion

Epidemiological characteristics of infectious diseases have changed dramatically requiring necessitate health care workers to understand the status of these diseases, especially for AIDS and other pandemics disease^[9–11]^.Efforts and investments have been made globally to strengthen the national HIV monitoring and evaluation (M&E) capacity^[12]^.Global priorities to end the HIV epidemic, such as achieving UNAIDS 90-90-90 targets, now focus on treatment as prevention and rapidly closing gaps in HIV prevention and care continues ^[13]^.Due to the wide use of highly active retroviral therapy and effective prevention measures, the AIDS epidemic had been effectively curbed, and the number of new HIV-1 infection had been declining^[14]^. Nevertheless, the HIV infection continued to be a serious threat to human health, particularly in developing countries. The first case of HIV-1 local infection was identified in the Dehong prefecture of Yunnan, China in 1989 ^[15]^. Since then, the number of HIV-1 infection has grown rapidly in China, which were also spread across the country^[16]^. On the basis of official statistics, the number of people living with HIV-1 was 780,000 in China at the end of 2011 ^[17]^. ^[18]^. Shanghai has the largest cross-border population entering China from countries around the world. Cross-border travel is important for culture and commerce but also leads to the spread of infectious diseases among countries. This was the first study which reported the HIV-1 infection rate and their characteristics among entry travelers over a long time span (2005-2016), among a large population (50830 cases) in Shanghai, China.

From 2005 to 2016, there were 50830 international travellers entering Shanghai who underwent physical examination in Shanghai International Healthcare Center. In this crossing border population, a total of 245 (0.48%) travelers were determined as HIV-1-positive. HIV incidence rate among active duty entry travelers in Shanghai peaked in 2014 and dropped to a stable but lower rate than observed in the USA and other underdeveloped countries ^[19–21]^. Overall, we found that the HIV-1 infection rate among entry travelers in Shanghai did not show an increasing tendency over these years, contributed by global HIV / Acquired Immunodeficiency Syndrome (AIDS) prevention and control efforts, such as promoting condoms in Southeast Asian countries and implementing the Asian regional AIDS project.

As a port health and quarantine organ, Shanghai International Travel Healthcare Center identifies AIDS patients and infection, mainly through the entry and exit personnel health examination. After the patients or the infected persons were found, the epidemic situations were reported through the CDC epidemic reporting system, and the related data were handed over to the local health department. In recent years, the Shanghai International Travel Healthcare Center has strengthened the follow-up supervision of the entry of HIV positive and AIDS patients, and established foreign liaison files for foreign HIV infected persons. Health education for the HIV infected persons was carried out to provide information about disease diagnosis, treatment and intervention for high risk behavior. With the local disease control departments, medical institutions, foreign embassies and other information interoperability, joint cooperation, explore overseas HIV infection health counseling and treatment measures. AIDS patients who do not want to return to their home country need to have relevant health consultation, and should be treated by designated medical institutions with paid medical treatment. Using the existing conditions, according to local conditions, the entry-exit personnel physical examination site could be set up as a consulting base. We tried to make full use of newspapers, magazines, radio, television and other mass media to answer typical questions;to establish International Travel Health Advisory contact, by using of websites, WeChat, telephone and other ways to answer questions raised by travelers on AIDS prevention and control work. With the local health administration, Family Planning Commission, Red Cross and other departments, through public welfare performances, on-site distribution brochures, free distribution of condoms and other means were undergoing.

In the last decade, we see a dramatic change in the demographic structure of the population of people living with HIV (PLWH)^[22]^. This study found that HIV-1 infection of entry travelers in Shanghai also had certain demographic characteristics. From the perspective of age-specific HIV-1 prevalence, the major age population of HIV-1 infected travelers was 18-40 years (73.90%). Overall, an estimated 5 million young people aged 15–24 were living with HIV in 2009 and accounted for 41% (about 890,000 cases) of new HIV infections globally ^[23,24]^. HIV is disproportionately afflicting young people worldwide, especially in the developing countries ^[25–27]^. Despite global commitment and ongoing prevention efforts, young people continue to demonstrate a sustained high risk for HIV ^[28]^. The similar trend was also found in our study. According to the diagnostic tests, those with positive results were divided into three groups,(more than 40 years old, 31 to 40 years old, less than 30 years old), working men aged 31 to 40 had the highest incidence whereas working women had a very low incidence, near zero. Our results are consistent with a previous study by Wang et.al, which also showed that travelers aged 21–30 and 31–40 were the most commonly infected individuals among entry travelers in Yunnan Province ^[2]^.

Demographic characteristics were also reflected in the marital status, working condition and education level. Travelers who were unemployed two years before detection constitute 15.51% of those that were tested positive while those who were employed constitute 53.88%. This result may be related to the characteristics of the total population. In addition, HIV-1 infection was more frequently detected among individuals with certain occupations such as businessman and entertainers. Furthermore, from a marriage point of view, unmarried (single or other marital status) applicants had higher proportion of infection compared to those married group(54.70%vs18.37%). In general, people with better education and better cognitive ability have healthier behaviors and therefore better health outcomes ^[31]^.Nevertheless, our study found that most of the infected people were highly educated, which were related to the characteristics of the applicants, and that most of the international travelers have high education level. The entry travelers are divided into high-income countries, middle-income countries, and low-income countries based on the host country ^[8]^. As expected, the data showed that more positive cases in this study were from middle-income or low-income countries.

From the view of transmission, our data suggested that sexual contact was the major high risk behavior, especially in those men who have sex with men (MSM). In the meantime, a study in the US reported the most common HIV transmission route had changed to homosexual contact in men, while injection drug users (IDU) was ranked the third^[32]^.Since sexual contact was identified as a major risk behavior in our research crowd, we deemed that the major HIV infection route had been shifted from intravenous drug use to sexual contact. Therefore, it suggested that more stringent surveillance of HIV-1 and other infectious pathogens in this population should be conducted, especially since the number of international travelers to Shanghai is increasing. Monitoring of HIV-1 infections in high-risk populations is particularly important for preventing the spread of HIV-1 infection across the border. This study found that those who did not use condom accounted for a large proportion in the positive cases. Inconsistent condom use and several risk-taking behaviors were also reported among young people in United States (African-Americans aged 18–21) and Uganda ^[33]^. Other studies also showed an increased rate of high-risk behaviors for HIV infection, particularly in unprotected sexual behavior ^[34–38]^. Insufficient HIV-related knowledge and low self-awareness of risk might be associated with an increasing number of AIDS. Thus, promotion and acceptance of integrated sex education incorporating accurate and age-appropriate information on HIV appears to be the need of the hour in order to interrupt the spread of HIV.

## Strengths and Limitations

Although the HIV-1 prevalence among these recruited 50830 international travelers who entered China at Shanghai port were described, there are still some limitations that need to be addressed. Since this was a record-based study, the records of some applicants were incomplete, preventing us from analyzing some variables. The face-to-face interviews were only conducted in the HIV population, and the characteristics of HIV negative population could not be obtained. Therefore we were not able to compare the demographic characteristics of HIV positive versus HIV negative cases. Furthermore, the sample may not be representative of the total population of entry travelers in China or other countries. At present, most foreigners in China are short-term visitors.Although some of them go back and forth many times, the time of staying in China is not more than one year each time, and the mobility of personnel is relatively large. AIDS monitoring for foreigners in China only applies to the foreign applicants who apply for residence in China, and there is no effective monitoring measures and methods for most of the short-term immigrants.

According to the relevant provisions, entry foreigners holding negative HIV antibody test results from overseas public hospitals can be exempted from inspection. However, some of these individuals’ HIV antibody test spot checks were found positive. At the same time, the public security, inspection and quarantine, education, human resources and social security departments found that some foreign health inspection materials provided by foreigners were incomplete or fraudulent. This part of the population can be the potential spread of AIDS population, their declaration of consciousness was very weak, and effective supervision of them was more difficult. However, the present study is one of the few similar studies that have been conducted in China. Although there were limitations and incomplete information, we still believed that this study could help us to understand the dynamics of HIV transmission among international travelers to some extent. The findings of this study also highlighted the information gap in this area.

## Conclusion

Our data from Shanghai International Travel Healthcare Center covering more than a decade, suggested a target for HIV/AIDS prevention and control efforts for international travelers. It was crucial to identify high-risk groups in which majority populations were infected, in order to make appropriate and targeted preventive efforts.

## Contributors

Jia Qin, Ru Zhou, Jing Xia, Weixin Wang, Jun Pan, Jiahong Pan, Xuan Zhou and Qi Zhang collaborated in the design of the study. Jia Qin and Ru Zhou contributed equally to this work, they wrote the initial drafts of this report.. All authors reviewed and edited the report and all take responsibility for its integrity and the accuracy of the analysis.

## Acknowledgements

We would like to thank all the participants of this study and all staff of HIV/AIDS case reporting system in Shanghai International Healthcare Center.

